# Signaling protein abundance modulates the strength of the Spindle Assembly Checkpoint

**DOI:** 10.1101/2022.05.10.491369

**Authors:** Chu Chen, Lauren Humphrey, Soubhagyalaxmi Jema, Shriya Karmarkar, Frank Ferrari, Ajit P. Joglekar

**Affiliations:** Department of Cell and Developmental Biology, University of Michigan Medical School, Ann Arbor, Michigan 48109; Department of Biophysics, University of Michigan, Ann Arbor, Michigan 48109; Department of Molecular Genetics of Ageing, Max Planck Institute for Biology of Ageing, Cologne, Germany 50931

## Abstract

During mitosis, unattached kinetochores in a dividing cell signal to the Spindle Assembly Checkpoint to delay anaphase onset and prevent chromosome missegregation ^1–4^. The signaling activity of these kinetochores and the likelihood of chromosome missegregation both depend on the amount of SAC signaling proteins that each kinetochore recruits ^5–8^. Therefore, factors that control SAC protein recruitment to signaling kinetochores must be thoroughly understood. Phosphoregulation of kinetochore and SAC signaling proteins emerging from the concerted action of many kinases and phosphatases is a major determinant of SAC protein recruitment to signaling kinetochores ^9^. Whether the abundance of SAC proteins also influences their recruitment and signaling activity at human kinetochores has not been studied ^8, 10^. Here, we reveal that the low cellular abundance of the SAC signaling protein Bub1 limits kinetochore recruitment of Bub1 and BubR1 and reduces the SAC signaling activity of the kinetochore. Conversely, Bub1 overexpression results in higher protein recruitment and SAC activity producing longer delays in anaphase onset. We also find that the number of SAC proteins recruited by a signaling kinetochore is inversely correlated with the total number of signaling kinetochores in the cell. This correlation likely arises from the competition among the signaling kinetochores to recruit from a limited pool of signaling proteins. The inverse correlation between the number of signaling kinetochores in the cell and the signaling activity of individual kinetochores may allow the dividing cell to prevent the large number of signaling kinetochores in prophase from generating an unnecessarily large signal, while enabling the last unaligned kinetochore to signal at the maximum possible strength.

## Results and Discussion

### The stoichiometry of SAC proteins at signaling kinetochores depends on the number of signaling kinetochores in the cell

We set two interrelated goals for this study. First, we wanted quantify the steady-state stoichiometry of key SAC proteins in human kinetochores to directly view the signaling activity of individual kinetochores ^6^. This has been accomplished in fungi ^8, 10^, but the data in vertebrates remains indirect and fragmentary ^5, 11^. Therefore, we quantified the recruitment of three SAC proteins representing the three layers of the well-defined signaling cascade that generates the ‘Mitotic Checkpoint Complex’ (MCC, Figure 1A) ^5, 12–21^. To accomplish this, we used three genome-edited HeLa-A12 cell lines wherein either Bub1, BubR1, or Mad1 was fused with mNeonGreen (mNG) ^22, 23^. These HeLa cell lines are only partially genome-edited; fully edited cell lines could not be obtained because of low efficiency of CRISPR-Cas9-mediated knock-ins and the pseudo- tetraploid nature of HeLa cells. Therefore, we also quantified the relative amounts of labeled and unlabeled proteins in the three cell lines using quantitative immunoblot analysis on whole-cell lysates of mitotic cells (Figure S1A-B). This analysis defined the labeled-to-unlabeled protein ratio for each of the three cell lines: ∼40% for Bub1 and ∼70% for both BubR1 and Mad1 (assuming similar transfer efficiencies for the labeled and unlabeled protein bands, Figure S1C). We used these ratios to estimate the total protein recruitment per kinetochore from the average signal for the mNG-tagged protein per kinetochore. It should be noted that these fusions do not detectably affect SAC signaling (shown later in Figure 4).

**Figure 1.**
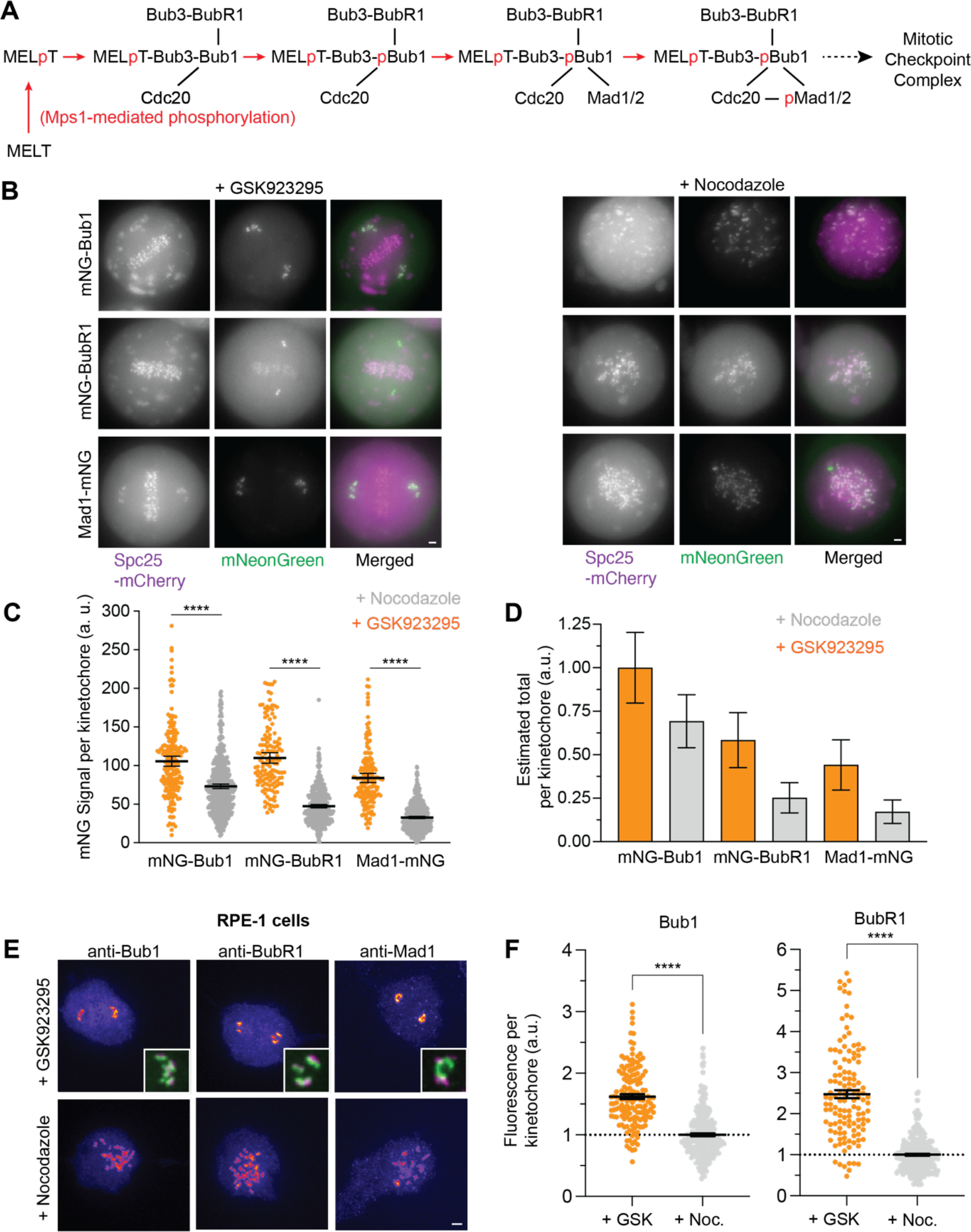
The SAC protein recruitment at signaling kinetochores depends on the number of signaling kinetochores in the cell. **(A)** A simplified schematic of the MELT motif-mediated recruitment of SAC proteins to the kinetochore. Red arrows signify Mps1-mediated phosphorylation, black lines indicate protein- protein interactions. The RZZ pathway, which is activated by Bub1 ^29, 35, 40^, is not shown. **(B)** Representative fluorescence micrographs of mitotic HeLa cells expressing the indicated proteins treated with GSK923295 (left) and nocodazole (right; scale bar ∼ 1.6 μm) **(C)** Quantification of the fluorescence signal per signaling kinetochore. Horizontal lines display mean ± 95% confidence intervals (n = 200, 142, 160 in GSK923295-treated cells and 562, 446, and 658 in nocodazole-treated cells for Bub1, BubR1, and Mad1 respectively, observations pooled from ≥ 2 technical repeats). Right: Spc25-mNG signal per kinetochore quantified from images acquired under identical imaging conditions (n = 178). Welch’s *t*-test was used to determine whether the sample means are significantly different. In all the figures in this study, **** indicates *p* < 0.0001, *** indicates 0.0001< *p* < 0.001, ** indicates 0.001 < *p* < 0.01, * indicates 0.01 < *p* < 0.05, and n.s. (not significant) indicates *p* > 0.05. **(D)** Estimation of total protein recruited per kinetochore based on the fluorescence measurements shown in C and immunoblot quantification of labeled to unlabeled fraction for each protein shown in Figure S1. **(E)** Representative immune-fluorescence micrographs (pseudo-colored as heatmaps) of RPE-1 cells. Insets show magnified regions of interest around unaligned chromosomes with antibodies labeling SAC proteins pseudo colored as green and antibodies labeling centromeres pseudo colored as magenta. Notice in the case of Mad1 a ring-like structure in the polar region presumably representing the Mad1 protein aggregated by its dynein-mediated transport from the kinetochore. This structure prevented us from quantifying the amount of kinetochore localized Mad1. Scale bar ∼ 2.44 μm. **(F)** Quantitation of immunofluorescence per kinetochore for Bub1 and BubR1 (each dot represents a single kinetochore; mean ± SEM is overlaid for each group. For Bub1, n = 255 (GSK923235-treated) and 157 kinetochores (nocodazole-treated) from two technical repeats. For BubR1, n = 125 (GSK923235-treated) and 288 kinetochores (nocodazole-treated) from two technical repeats. Welch’s *t*-test used to compare sample means.

For our second goal, we studied whether the number of signaling kinetochores in the cell affects the signaling activity of individual kinetochores. In budding yeast, the recruitment of SAC proteins per kinetochore is higher in cells that contain fewer signaling kinetochores, likely because of the limited abundance of SAC proteins ^10^. Whether this factor influences the signaling activity of human kinetochores is unknown. Therefore, we observed SAC protein recruitment per kinetochore in mitotically arrested cells containing two distinctly different numbers of signaling kinetochores. To obtain cells with nearly all kinetochores signaling, we released G1/S arrested HeLa cells into the cell cycle and then arrested them in mitosis using 330 nM nocodazole to depolymerize microtubules (Figure 1B, right). These cells serve as a model for late prophase. To obtain mitotic cells with a much smaller number of signaling kinetochores, we released G1/S arrested HeLa cells into the cell cycle and treated them with 236 nM GSK923295, a small molecule inhibitor of the mitotic kinesin CENP-E, 1.5 hours prior to imaging ^24^. These cells model late prometaphase, because most of the chromosomes align at the metaphase plate, but a smaller, variable number of chromosomes become stranded at the spindle poles. The unattached or laterally attached kinetochores of these chromosomes activate the SAC (Figure 1B, left) ^25, 26^. Relying on visual inspection, we selected only those cells containing ∼ 10 or less polar chromosomes for analysis.

The quantitative comparison of mNG-labeled protein recruitment revealed that the recruitment of all three proteins per kinetochore was significantly higher in GSK923295-treated compared to nocodazole-treated cells (Figure 1C). We combined these measurements with the quantitative immunoblotting data to estimate the total (labeled + unlabeled) protein amount to understand how BubR1 and Mad1 recruitment correlates with Bub1 recruitment after the two treatments (Figure 1D). We normalized total protein amounts with the amount of total Bub1 per kinetochore in GSK923295-treated cells ^13, 27–32^. In GSK923295-treated cells, BubR1 and Mad1 per kinetochore was ∼ 60 and 45% respectively of total Bub1 per kinetochore. In nocodazole-treated cells, the BubR1 and Mad1 amount per kinetochore was significantly lower ∼ 40% and 25% respectively of total Bub1. Thus, the overall protein recruitment per kinetochore was higher in GSK923295- treated cells compared to nocodazole-treated cells: 1.45-fold in case of Bub1, 2.3-fold and 2.6- fold for BubR1 and Mad1 respectively (Figure 1D). Thus, the number of SAC proteins recruited per kinetochore in HeLa cells is inversely correlated with the number of signaling kinetochores in the cell, consistent with observations in budding yeast ^10^.

We used a fluorescence standard to estimate the copy number of each SAC protein recruited per kinetochore. It is known that: (a) the human kinetochore contains ∼ 250 molecules of the Ndc80 complex and (b) the stoichiometry between Ndc80 and Knl1 is 3:2 ^33^. Therefore, under identical imaging conditions we imaged HeLa cells exogenously overexpressing Spc25-mNG (Spc25 is an Ndc80 complex subunit) and treated with siRNA targeting the endogenous Spc25 to quantify the mNG signal per kinetochore in metaphase cells (Figure S1D). Using the above-mentioned parameters, we converted the estimated total amount of each protein per kinetochore into the number of molecules of the protein recruited per Knl1 molecule in HeLa cells. These calculations indicate that each Knl1 molecule recruits 5 ± 0.9 Bub1 molecules, 3 ± 0.7 BubR1 molecules, and 2 ± 0.7 Mad1 molecules in GSK923295-treated cells. The numbers are lower in nocodazole-treated cells: 3 ± 0.7 Bub1 molecules, and 1 ± 0.4 molecules of BubR1 and Mad1 each per Knl1. The number of Bub1 molecules per kinetochore in nocodazole-treated cells is slightly lower than the previously reported number of 4-6 molecules ^5, 12, 13^. A more recent study found that Bub1 recruitment per kinetochore was not significantly different in cells treated with nocodazole- and GSK923295 ^34^. However, this difference could arise from the lack of filtering based on the number of signaling chromosomes per cell.

Mad1 is recruited to metazoan kinetochores by Bub1 and the RZZ complex ^35–37^. To quantify the contribution of the RZZ complex, we depleted *ZW10* using RNA interference (RNAi) and then quantified SAC protein recruitment as before. *ZW10* siRNA treatment lowered Bub1 and BubR1 recruitment by ∼ 25%, as noted by others ^38^. As expected, Mad1 recruitment was significantly lower in both GSK923295 and nocodazole-treated cells (Figure S1). The number of Mad1 molecules recruited per Bub1 molecule was ∼ 50% lower revealing the contribution of the RZZ under these conditions. Although the number of Bub1 and BubR1 molecules per kinetochore was also reduced, the two proteins were recruited in approximately equal amounts.

To test whether the inverse correlation is specific to HeLa cells, we examined Bub1 and BubR1 recruitment per kinetochore in hTERT-RPE1 cells. We released RPE-1 cells synchronized in G1/S into media containing either nocodazole or GSK923295, and measured the total amounts of Bub1, BubR1 recruited per kinetochore by immunofluorescence (Figure 1E). These measurements show that the Bub1 amount per kinetochore in GSK923295-treated RPE-1 cells as ∼1.6-fold higher the Bub1 amount in nocodazole-treated cells; BubR1 amount was ∼2.5-fold higher (Figure 1F). We did not quantify the amounts of Mad1 per kinetochore because a ring-like pool of Mad1 proximal to the spindle poles, especially in GSK923295-treated cells (see inset in Figure 1E), prevented accurate quantification. These data demonstrate that the inverse correlation between the amount of Bub1 and BubR1 per kinetochore and the number of signaling kinetochores is not specific to HeLa or budding yeast cells. In fact, Bub1 and BubR1 show a similar degree of enrichment at RPE-1 and HeLa kinetochores in GSK923295-treated cells.

### Mps1-mediated phosphorylation within signaling kinetochores is the same in nocodazole- and GSK923295-treated cells

Changes in the Mps1-mediated phosphoregulation of the kinetochore in GSK923295- and nocodazole-treated cells may change SAC protein recruitment. Indeed, a prior study found that the net Mps1-mediated phosphorylation decreased rather than increasing in GSK923295-treated cells when compared to nocodazole-treated cells suggesting that SAC protein recruitment should decrease in GSK923295-treated cells ^34^. Therefore, we quantified the net Mps1 kinase activity within the kinetochore using a recently developed FRET-based sensor (MPS1sen-KT) ^34, 39^. We modified this sensor to use mNeonGreen/mScarlet-I as the acceptor/donor combination (Figure S2A). Since the acceptor and the donor has a fixed 1:1 stoichiometry in the sensor, we used the ratio between the green channel readout and the FRET channel readout (without correcting for the cross-excitation of the acceptor fluorescence and the bleed-through of donor fluorescence) as the normalized measurement of the FRET efficiency, which is positively correlated with the net phosphorylation (Figure S2B-D).

We confirmed that MPS1sen-KT recruited to unaligned kinetochores in both nocodazole- and GSK923295- treated cells had lower FRET efficiencies (higher activities of Mps1) compared to kinetochores aligned at metaphase plates in GSK923295-treated cells (Figure S2D). Importantly, we did not detect any difference in the net Mps1-mediated phosphorylation within unaligned kinetochores in nocodazole- and GSK923295-treated cells irrespective of the drug concentration used (Figure S2D). This divergence which may arise from different properties of the two cell lines as was found in a panel of colorectal cancer cell lines ^34^. In either case, the increased kinetochore recruitment of SAC proteins in GSK929395-treated cells does not stem from increased Mps1- mediated phosphorylation within the kinetochore.

### The number of MELT motifs per Knl1 controls Bub1, BubR1, and Mad1 recruitment to signaling kinetochores

We next studied how the number of MELT motifs per Knl1 affects the SAC protein recruitment. We knocked down endogenous Knl1 using RNAi and replaced it with Knl1^Δ^-M3 or Knl1^Δ^-M3-M3, which are mScarlet-I-tagged recombinant Knl1 versions containing three or six MELT motifs: (Figure 2A) ^5^. As before, we quantified Bub1, BubR1, and Mad1 signals at individual kinetochores in cells treated with either GSK923295 or nocodazole (Figure 2B). For comparing the amounts of proteins recruited under these conditions, we normalized all localized fluorescent intensities by the average fluorescence signal per kinetochore of each protein in nocodazole-treated cells expressing Knl1^Δ^-M3. As before, we used the labeled-to-unlabeled protein ratios to estimate the total amount of protein per kinetochore (Figure 2C).

**Figure 2.**
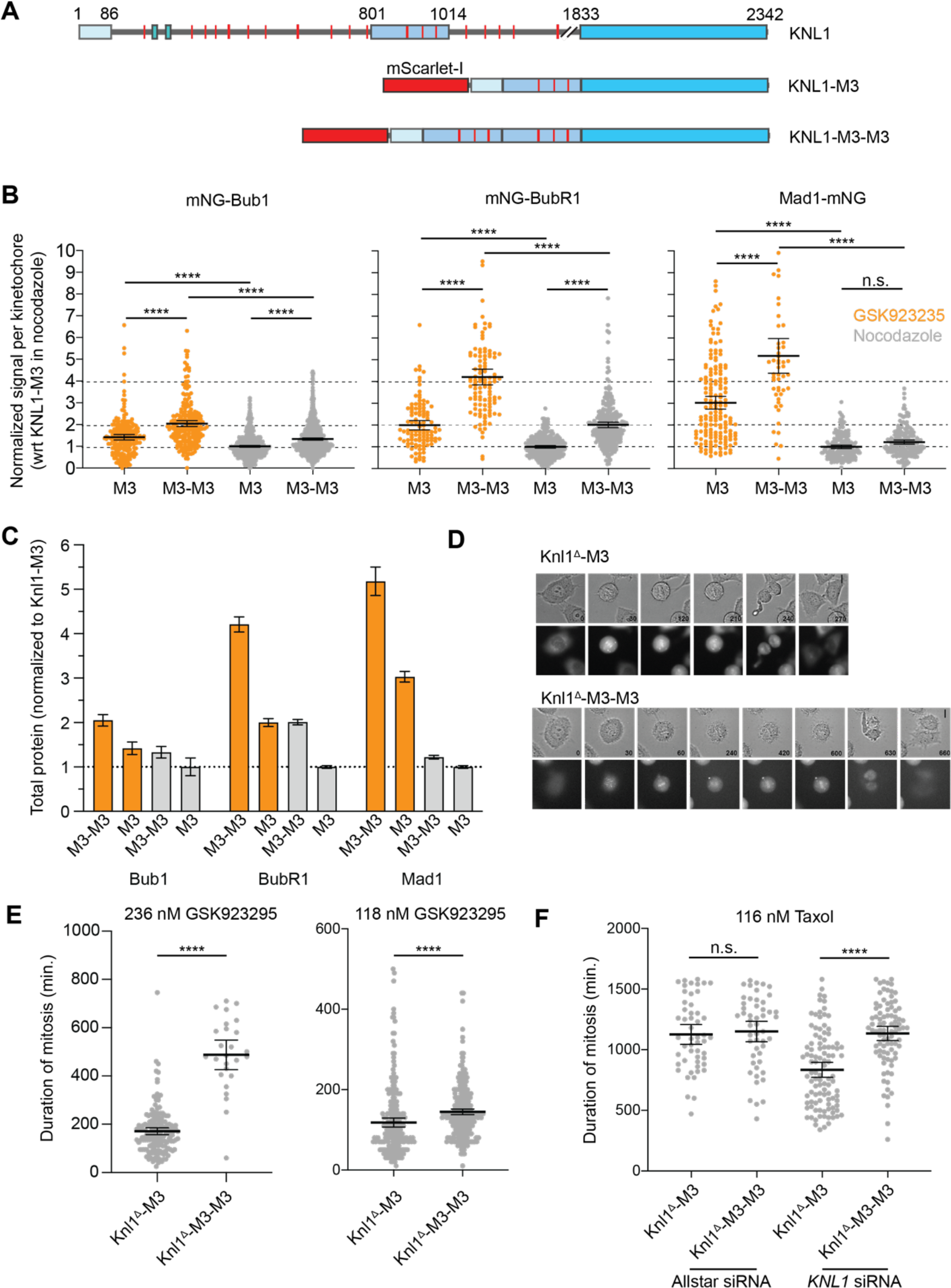
The number of MELT motifs in Knl1 affects SAC protein recruitment to the kinetochore and SAC signaling strength. **(A)** Schematics of the full-length, endogenous Knl1 and the recombinant Knl1^Δ^-M3 and Knl1^Δ^- M3-M3 versions. The expression of Knl1^Δ^-M3 (or Knl1^Δ^-M3-M3) is under the regulation of a constitutive *EF1a* promoter. **(B)** Quantification of the fluorescence signal per signaling kinetochore for the indicated protein. In each case, the signals were normalized using the mean signal for each protein measured in nocodazole-treated cells expressing Knl1^Δ^-M3 (mNG-Bub1: n =844 and 602 for Knl1^Δ^-M3-M3 and Knl1^Δ^-M3 respectively in nocodazole-treated cells and 222 and 120 for Knl1^Δ^- M3-M3 and Knl1^Δ^-M3 respectively in GSK923295-treated cells, pooled from ≥ 2 technical repeats; mNG-BubR1: n = 278 and 272 for Knl1^Δ^-M3-M3 and Knl1^Δ^-M3 respectively in nocodazole-treated cells and 110 and 107 for Knl1^Δ^-M3-M3 and Knl1^Δ^-M3 respectively in GSK923295-treated cells, pooled from ≥ 2 technical repeats; Mad1-mNG: n = 182 and 184 for Knl1^Δ^-M3-M3 and Knl1^Δ^-M3 respectively in nocodazole-treated cells and 52 and 166 for Knl1^Δ^-M3-M3 and Knl1^Δ^-M3 respectively in GSK923295-treated cells, pooled from ≥ 2 technical repeats). **(C)** Estimation of total protein amount per kinetochore relative to the total Bub1 amount per kinetochore in drug-treated cells (colors same as in as in B). **(D)** Representative micrographs show mitotic progression of cells with either Knl1^Δ^-M3 (top) or Knl1^Δ^-M3-M3 (bottom, scale bar: 10 μm). Time stamps are in minutes. **(E-F)** Duration of mitosis for cells treated with the indicated concentration of GSK923295 and Taxol (left: n = 162 and 26 for Knl1^Δ^-M3 and Knl1^Δ^-M3-M3 respectively from two technical repeats; right: n = 235 and 339 for Knl1^Δ^-M3 and Knl1^Δ^-M3-M3 respectively, n = 51 and 52 for Allstar siRNA + taxol treatment and n = 102 and 88 for KNL1 siRNA + taxol treatment for Knl1^Δ^-M3 and Knl1^Δ^- M3-M3 respectively). Welch’s *t*-test was used to compare sample means.

In the GSK923295-treted cells, Knl1^Δ^-M3 recruited ∼ 1.4-times higher Bub1 per kinetochore compared to nocodazole-treated cells. Knl1^Δ^-M3-M3 recruited more Bub1 than Knl1^Δ^-M3 under both conditions. Importantly, Knl1^Δ^-M3-M3 recruited 2-fold higher Bub1 than Knl1^Δ^-M3 in GSK923295-treated cells consistent with its 2-fold higher number of MELT motifs. These data again highlight that the recruitment of SAC proteins per kinetochore is higher in cells containing small numbers of signaling kinetochores. BubR1 and Mad1 recruitment showed similar trends.

Importantly, the total BubR1 recruitment per Knl1^Δ^-M3-M3 was 2-fold than Knl1^Δ^-M3 under both conditions. BubR1 amount per Knl1^Δ^-M3 was 2-fold higher in GSK923295-treated cells compared to nocodazole-treated cells, which is consistent with the increase in Bub1 recruitment by Knl1^Δ^- M3 under the two conditions. Interestingly, total Mad1 recruitment by Knl1^Δ^-M3-M3 and Knl1^Δ^-M3 was similar in nocodazole-treated cells but increased by 5-fold in GSK923295-treated cells. The kinetochore recruitment of the two recombinant Knl1’s was very similar and cannot explain the observed trends in SAC protein recruitment (Figure S3A).

As before, we combined these data with the immunoblotting quantitation to estimate the number of molecules per Knl1 (Figure S3B). Knl1^Δ^-M3-M3 cells recruits ∼ 5 ± 0.3 Bub1 molecules, but only 3 ± 0.3 molecules in nocodazole-treated cells. These numbers are very similar to the numbers of Bub1 molecules recruited by wild-type Knl1 under the two conditions (5 ± 0.9 and 3 ± 0.7 molecules respectively, see Figure S1), indicating that our estimates are internally consistent. A prior study reported that HeLa kinetochores recruit 4-6 Bub1 molecules per kinetochore in nocodazole-treated cells ^5^, whereas we find that there are 3 Bub1 molecules per Knl1 in nocodazole-treated cells and 5 per kinetochore in GSK923295-treated cells. The reasons for this discrepancy remain unclear.

These data provide several insights. First, they confirm that the number of MELT motifs per Knl1 strongly influences Bub1, BubR1, and Mad1 recruitment. Second, in the case of Bub1 and Mad1 this influence is detected only in GSK923295-treated cells containing small numbers of signaling kinetochores. Third, Knl1^Δ^-M3-M3 recruits ∼2-fold more BubR1 than Knl1^Δ^-M3 under both conditions. Increased BubR1 recruitment to the kinetochore is likely to be functionally significant because it promotes SAC signaling ^14, 23^. Mad1 recruitment is disproportionately high GSK923295- treated cells, possibly because of changes in the contribution of the RZZ pathway to Mad1 recruitment ^40^. Given the catalytic role of Mad1 in the rate-limiting step in the SAC signaling cascade, the higher Mad1 recruitment may also increase the signaling activity of these kinetochores. Therefore, we next tested whether the higher SAC protein recruitment by Knl1^Δ^- M3-M3 translates into a stronger SAC and longer mitotic arrest durations.

### The number of MELT motifs per Knl1 affects the SAC signaling strength

Previous studies concluded that the number of MELT motifs per Knl1 does not significantly influence the duration of the SAC-mediated mitotic arrest unless the MELT motifs are weak in their activity ^5, 13^. However, these studies characterized SAC signaling only in cells treated with high doses of nocodazole. Our quantification shows that in the nocodazole-treated cells, kinetochores containing Knl1^Δ^-M3-M3 recruit only modestly larger amounts of Bub1 and Mad1 compared to kinetochores containing Knl1^Δ^-M3. Therefore, the SAC signaling activity of both types of kinetochores may have been similar in nocodazole-treated cells. A further complicating factor is that one of the studies partially inhibited Mps1 during the experiment, which may have masked differences in SAC signaling due to the two Knl1 versions ^5^.

Because of these considerations, we re-evaluated the strength of SAC signaling mediated by Knl1^Δ^-M3 and Knl1^Δ^-M3-M3 specifically in GSK923295-treated cells. We used time-lapse imaging to monitor mitotic progression in cells treated with different concentrations of GSK923295 to achieve different degrees of CENP-E inhibition resulting in different durations of mitotic arrests. In all cases, cells expressing Knl1^Δ^-M3-M3 arrested for significantly longer durations compared to cells expressing Knl1^Δ^-M3 (Figure 2D-E). The same trend was observed when these cells were treated with Taxol (Figure 2F). Thus, the number of MELT motifs in Knl1 can determine the duration of checkpoint-induced mitotic delays in cells wherein a few signaling kinetochores sustain the SAC. When combined with the results described in the previous section, this finding implies that the higher recruitment of Bub1, BubR1, and Mad1 per kinetochore is correlated with a higher SAC activity, and this correlation is readily detected only in when the mitotic cells contain small numbers of signaling kinetochores ^7^.

### Bub1 over-expression increases recruitment of SAC proteins to the kinetochore

A key question then what limits SAC protein recruitment to the kinetochore especially in nocodazole-treated cells. Changes in phosphoregulation cannot explain the differences in SAC protein recruitment under the two conditions studied here. One responsible factor may be the low abundance of SAC proteins, especially Bub1, which is responsible for recruiting BubR1 and Mad1. Bub1 overexpression results in higher Bub1 and Mad1 recruitment to the budding yeast kinetochore ^10^. Mad1 recruitment may also be limited by its own low abundance compared to Bub1 and BubR1 ^23^. Therefore, to assess SAC protein abundance, we used Fluorescence Correlation Spectroscopy (FCS). We performed FCS measurements on mitotically cells arrested either using MG132 or nocodazole. These measurements identified Mad1 is the protein with the lowest abundance amongst the three SAC proteins. They also suggest that cytosolic concentration of Bub1 is modestly lower in nocodazole-treated compared to MG132-treated cells (Figure S3C).

To test whether higher Bub1 expression can lead to higher kinetochore recruitment of SAC proteins, we exogenously expressed mNG-Bub1 in HeLa cells using a doxycycline-inducible promoter (Figure S4A) and quantified Bub1, BubR1, and Mad1 recruitment per kinetochore in cells treated with GSK923295 using immunofluorescence. As expected, mNG-Bub1 localization at signaling kinetochores was detected only after doxycycline treatment (Figure 3A). To measure the total amount of Bub1 per kinetochore, we fixed and stained these cells with anti-Bub1 antibodies. Fluorescence quantification confirmed that signaling kinetochores in the doxycycline-treated cells recruited ∼35% more Bub1 than signaling kinetochores in untreated cells (Figure 3A, plot on the left). The kinetochores in the doxycycline-treated cells also recruited 26% more BubR1 (Figure 3A, middle plot), however, Mad1 amount per kinetochore did not change (Figure 3A, right plot). Thus, signaling kinetochores in HeLa cells can recruit more Bub1 and BubR1 if more Bub1 is made available. However, Mad1 recruitment does not increase likely because of the low Mad1 abundance, although behavior of the RZZ pathway will also strongly influence Mad1 recruitment.

**Figure 3.**
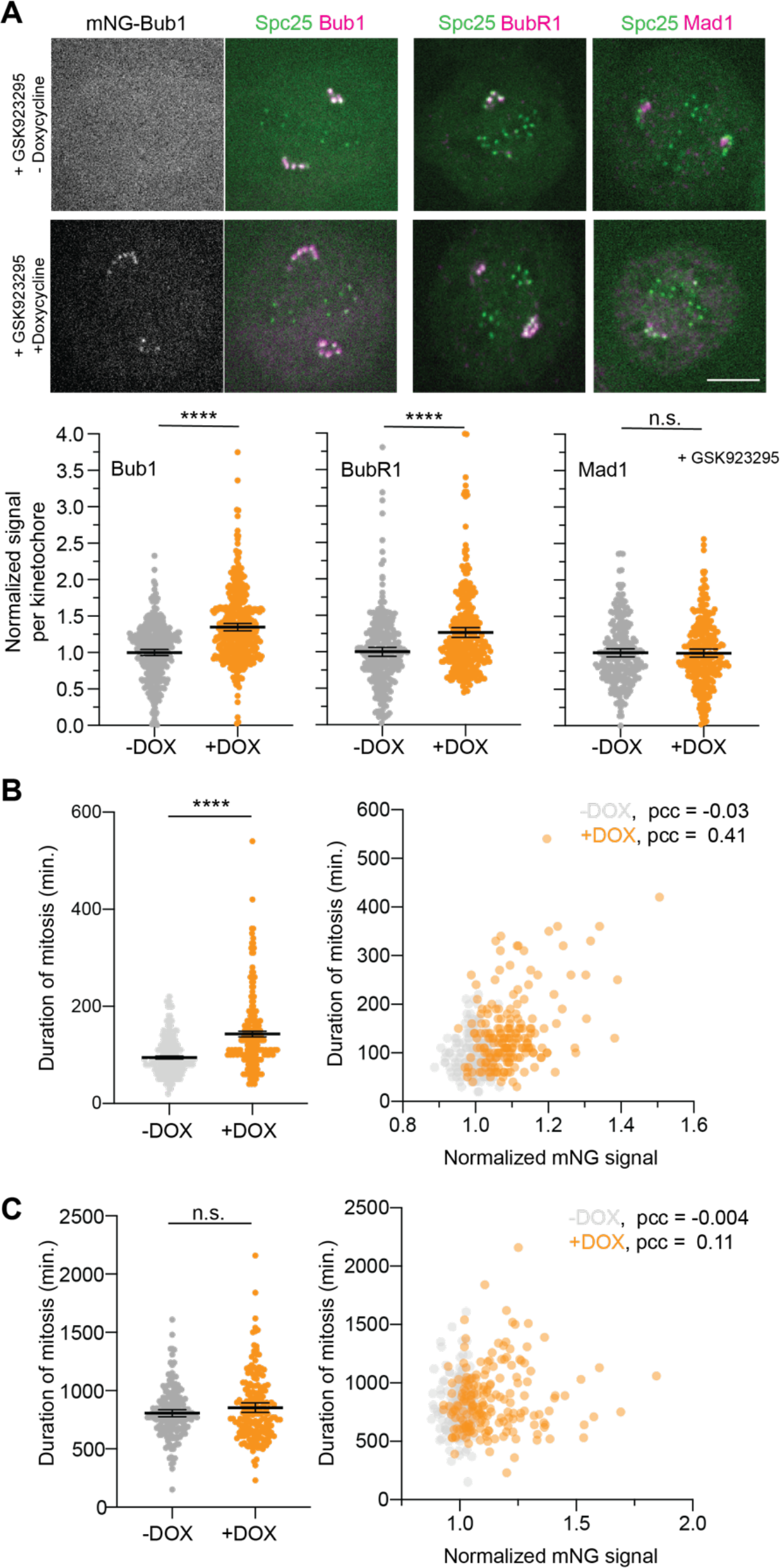
Bub1 over-expression results in higher Bub1 and BubR1 recruitment per kinetochore and prolongs the delay in anaphase onset. **(A)** Top panel: representative fluorescence micrographs of control (top) cells and cells treated with doxycycline to exogenously express mNG-Bub1. Scale bar ∼ 4.92 μm. Bottom panel: quantification of immunofluorescence signal per kinetochore for the indicated proteins in control cells and cells treated with doxycycline to induce exogenous mNG-Bub1 expression. Signal values were normalized with the average signal for the respective protein in doxycycline-untreated cells. (n = 344 and 390 for Bub1, 346 and 387 for BubR1, and 234 and 275 for Mad1 in untreated and doxycycline-treated cells respectively, observations pooled from 3 technical repeats). Welch’s *t*-test used to compare sample means. **(B)** Left: Duration of mitosis in control and doxycycline-treated cells in media containing 25- 37 nM GSK923295 (n = 195 and 187 for untreated and doxycycline-treated cells respectively, from 2 technical repeats; p < 0.0001 for doxycycline-treated and GSK923295 treated cells). Horizontal lines represent mean ± 95% confidence intervals on the mean. Right: Correlation between the duration of mitosis and the average cellular level of the exogenously expressed mNG-Bub1. mNG-Bub1 signal per cell was normalized with the average signal measured in control cells. Pearson’s correlation coefficient (pcc) for each data set is noted in the figure (*p =* 0.6222 and < 0.0001 respectively for untreated and doxycycline-treated cells respectively) **(C)** Left: Duration of mitosis in control and doxycycline-treated cells in media containing 330 nM nocodazole (n = 189 and 185 for untreated and doxycycline-treated cells respectively, from 2 technical repeats). Horizontal lines represent mean ± 95% confidence intervals on the mean. Right: Correlation between the duration of mitosis and the average cellular level of the exogenously expressed mNG-Bub1. mNG-Bub1 signal per cell was normalized with the average signal measured in control cells. Pearson’s correlation coefficient (pcc) for each data set is noted in the figure (*p =* 0.053 and 0.1309 respectively for untreated and doxycycline-treated cells respectively).

### Bub1 overexpression strengthens the SAC in the presence of a small numbers of signaling kinetochores

To test whether Bub1 overexpression results in longer delays in anaphase onset, we released either untreated or doxycycline treated, G1/S synchronized cells into media containing 18-35 nM GSK923295 and observed their mitotic progression (Figure 3B). We found that the doxycycline-treated cells arrested for ∼ 145 min on average, whereas the untreated cells arrested for ∼ 94 min under the same condition (Figure 3B, left, also see Supplementary Videos S1 and S2). To understand the correlation between the expression level of exogenous Bub1 and the mitotic duration, we plotted the mitotic duration against the cellular mNG signal in both cases (the fluorescence values were normalized to the average mNG fluorescence per cell from samples not treated with doxycycline, 3B right). As expected, there was no correlation between the cellular mNG signal and mitotic duration in untreated cells (gray circles, Pearson’s correlation coefficient displayed at the top right of the figure). The mitotic duration of doxycycline-treated cells was positively correlated with the mNG-Bub1 signal (orange circles in Figure 5B). It should be noted that the mNG-Bub1 over-expression did not affect the mitotic progression of normally dividing cells, and the degree of overexpression was also uncorrelated with the duration of mitosis (Figure S4B-C). Thus, the longer mitotic duration seen in GSK923295-treated cells arises from higher SAC signaling activity of the unaligned kinetochores.

We also examined whether Bub1 overexpression produces longer delays in anaphase onset in nocodazole-treated cells (Figure 3C, also see Supplementary Videos S3 and S4). Interestingly, the cellular mNG signal and the duration of mitosis did not statistically significant correlation in this case (Figure 3C). The likely explanation for this observation is that even with increased Bub1 recruitment, the large number of signaling kinetochores cannot recruit higher amounts of BubR1/MAD1 because of their depletion (implied by the reduced BubR1/MAD1 recruitment in the nocodazole-treated cells, Figure 1). Therefore, the Bub1 overexpression does not translate into more SAC signaling activity. In conclusion, increased Bub1 levels in HeLa cells increases Bub1 and BubR1 recruitment to the kinetochore and strengthens the SAC. Importantly, the effect on SAC strength is detected only in GSK923295-treated cells, which contain small numbers of signaling kinetochores.

### Partial depletion of Bub1 and BubR1 weakens the SAC in cells containing small numbers of signaling kinetochores

Mad1 abundance is lower than Bub1 abundance, which suggests that Mad1 availability should also limit the signaling activity of individual kinetochores (Figure S3C). Because Mad1 over- expression can weaken the SAC by depleting the Mad2 pool available for MCC formation, we used a different approach ^8, 41–43^. Rather than over-expressing Mad1, we partially knocked down Mad1-mNG using an siRNA against mNeonGreen in the genome-edited cell line heterozygous for Mad1-mNG, and then quantified the duration of SAC-induced mitotic delays. We also lowered mNG-Bub1 and mNG-BubR1 protein using the same strategy.

As expected, the mNG-tagged protein was significantly depleted by the mNeonGreen siRNA after two days, whereas the unlabeled protein was unaffected (Figure 4A). To study the effect of a mild depletion of an SAC protein on the signaling activity of the kinetochore, we treated the three cell lines with mNG siRNA for one day and observed the effects of protein depletion on mitotic duration in GSK923295- and nocodazole-treated cells. Quantification of mNG fluorescence signal from mitotic cells revealed that the mNG RNAi reduce the level of the mNG-tagged protein by ∼ 50% (Figure 4B). In GSK923295-treated cells, it also significantly affected the mitotic progression (Figure 4C, left). In the case of mNG-Bub1 and mNG-BubR1 depletion, the average duration of mitosis decreased from ∼ 1000 minutes to 300-500 minutes (Figure 4C). We also confirmed that the partial depletion of BubR1 in GSK923295-treated cells resulted in chromosome missegregation (Supplementary Videos S5 and S6). Mad1 depletion had a relatively minor effect on mitotic duration: the average time in mitosis decreased from ∼ 1000 minutes to 868 minutes (Figure 4C). Interestingly, in nocodazole-treated cells, the partial depletion of all three proteins did not significantly change the duration of mitosis (Figure 4C, right).

**Figure 4.**
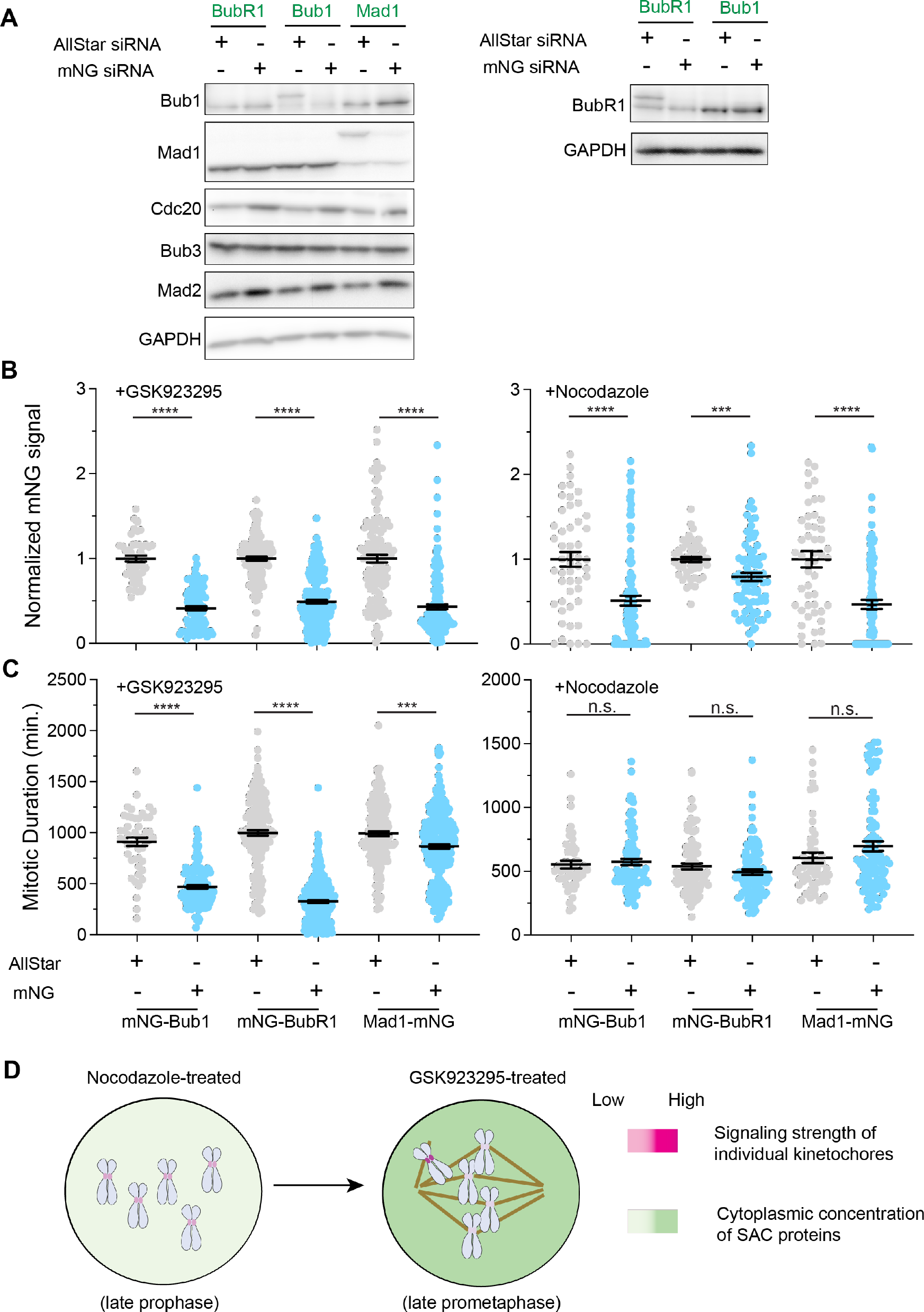
Partial depletion of Bub1 and BubR1, but not Mad1, weakens the signaling potential of the SAC. **(A)** Immunoblot analysis of total cell lysates shows that the mNG-tagged SAC protein is specifically targeted for RNAi after a 48-hour treatment with mNG siRNA. **(B)** Quantification of the mNG-signal per cell under the indicated conditions (For the experiment involving GSK923295 treatment, n = 50, 126, 130 for mNG-Bub1, mNG-BubR1, and Mad1-mNG cells respectively treated with Allstar siRNA; n = 100, 219, and 177 for mNG-Bub1, mNG-BubR1, and Mad1-mNG cells respectively treated with mNG siRNA; data collected in two technical repeats). For the experiment involving nocodazole treatment, n = 50, 50, and 50 for mNG-Bub1, mNG-BubR1, and Mad1-mNG cells respectively treated with Allstar siRNA; n = 100, 85, and 100 for mNG-Bub1, mNG-BubR1, and Mad1-mNG cells respectively treated with mNG siRNA; data collected from 2 technical repeats). Each dot represents a measurement from one cell, mean ± SEM overlaid. Welch’s *t*-test used to compare sample means. **(C)** Duration of mitosis for same cells as in B in media containing GSK923295 and nocodazole (for GSK923295-treated cells: *p* ≤ 0.0001 based on ordinary one-way ANOVA; for nocodazole- treated cells, *p* > 0.253 based on ordinary one-way ANOVA). Each dot represents a measurement from one cell, mean ± SEM overlaid. Pairwise comparisons of sample means performed using Tukey’s multiple comparison test. **(D)** Model depicts the combined effect of the number of signaling kinetochores per cell and the availability of free, cytoplasmic SAC proteins on the signaling strength of individual kinetochores.

Many prior studies have shown that the depletion of Bub1 in HeLa cells has only a minor effect on the duration of mitosis in nocodazole-treated cells ^35, 44–46^. Our results in nocodazole-treated cells are consistent with these findings. Strikingly, we find that in GSK923295-treated cells even a small (∼ 20-40% on average) reduction in Bub1 or BubR1 level results in a significant decrease in the duration of mitosis. This observation is consistent with our results involving Knl1^Δ^-M3 and Knl1^Δ^-M3-M3 (Figure 2) and suggests that higher SAC protein recruitment to the kinetochore results in higher SAC signaling activity. The likely reason why this is not seen in nocodazole-treated cells is that the combined activity of a large number of signaling kinetochores masks defects in the signaling activity of individual kinetochores ^7^.

In conclusion, HeLa and RPE-1 cells containing small numbers of signaling kinetochores recruit significantly higher amounts of Bub1, BubR1, and Mad1 per signaling kinetochore compared to cells containing many signaling kinetochores. The recruitment of all three proteins strongly correlates with the number of MELT motifs in Knl1. This correlation is readily detected in cells containing small numbers of signaling kinetochores; it cannot be detected in cells containing many signaling kinetochores likely because of a shortage of unbound SAC proteins. Importantly, the increased SAC protein recruitment per kinetochore, either due to the larger number of MELT motifs or Bub1 over-expression, results in longer delays in anaphase onset. Conversely, partial depletion of Bub1 and BubR1, but not Mad1, leads to shorter delays in anaphase onset in the presence of a small number of signaling kinetochores.

The inverse correlation between the recruitment level of SAC proteins at a signaling kinetochore and the total number of signaling kinetochores in a cell is likely to be biologically significant (Figure 4D). In human cells, the number of signaling kinetochores drops by almost two orders of magnitude: from ∼ 92 following nuclear envelop break-down to a few or just one in prometaphase. If the amount of MCC in the cell scales linearly with the number of signaling kinetochores, the nearly 100-fold drop in the number of kinetochores will result in a similar drop in the MCC creating the possibility of a premature anaphase onset. Pre-mitotic MCC generation at the nuclear envelop also buffers SAC signaling activity of the kinetochores ^47^. However, the strong decrease in the duration of mitotic arrest in cells expressing Knl1^Δ^-M3 or cells with partial Bub1 depletion suggests that the buffering mechanism cannot compensate for lower signaling activity of kinetochores. Instead, our data suggest that as the number of signaling kinetochores drops, the availability of SAC signaling proteins, especially Bub1, increases, and enables the remaining signaling kinetochores to recruit more SAC proteins. Consequently, the signaling strength of each kinetochore will become inversely correlated to the number of signaling kinetochores in the cell. Additionally, synergistic signaling by the Knl1 phosphodomain under this condition may further strengthen the signaling activity of these remaining kinetochores ^48^.

Our results also raise additional questions requiring further investigation. First, it remains unclear why kinetochores in HeLa cells and other eukaryotes use only a small fraction of their signaling strength. One possibility is that this modulation of the signaling strength of the kinetochore represents a compromise for balancing SAC signaling strength with its responsiveness to silencing mechanisms ^7^. Higher Bub1 expression may also have drawbacks in other aspects of mitosis and cell biology ^49^. Finally, these results bring into focus the contribution of perturbations or imbalance in the expression levels of SAC proteins on chromosome missegregation commonly seen in cancer cells ^50^.

## Materials and Methods

### Cell culture

HeLa A12 cells (a kind gift from the Makeyev lab) were grown in DMEM media supplemented with 10% fetal bovine serum (FBS), Pen-Strep, 25 mM HEPES, and 1x GlutaMAX at 37 °C and 5% CO2. 0.5-2 μg/ml Puromycin or 10 μg/ml Blasticidin was used as needed. To express recombinant proteins, we modified a bicistronic vector designed by the Makeyev lab as necessary ^51^. The modified vectors were transfected into HeLa A12 cells that have the *LoxP*-Blasticidin-*Lox2272* cassette engineered into their genome. Transformed cells were pooled and used for further experimentation. The methodology used to construct the genome-edited HeLa A12 cell lines has been described in detail in ^52^.

Imaging experiments were performed in Fluorobrite media supplemented with FBS, Pen-Strep, HEPES as noted above. G1/S synchronization was achieved using a double thymidine block procedure (2.5 mM thymidine). On the day of imaging, cells were released from the second block and treated with either nocodazole or GSK923295 ∼ 1.5 hours prior to imaging.

### Immunoblotting

For the quantitative immunoblot analysis shown in Figure S1, cells were synchronized using a single thymidine block (2.5 mM thymidine) for 24 hours. The cells were released into media containing 660 nM Nocodazole and harvested 15 hours post release. Anti-Bub1 (A300-373A, Bethyl Laboratories), anti-BubR1 (A300-386A, Bethyl Laboratories), and anti-Mad1 (ab184560, Abcam) primary antibodies were used to probe membranes at a dilution of 1:1000. Alexa Fluor 633-conjugated secondary antibodies (Goat anti-Mouse, Invitrogen A21052 or Goat anti-Rabbit, Invitrogen A21071 as appropriate) were used at a 1:15000 dilution.

For immunoblots in Figures 4A, S2A, and S4A: To acquire unsynchronized HeLa-A12 cells lysates, cells were either scraped off the dish surface or trypsinized. To acquired mitotic HeLa- A12 cells, cells were first synchronized in G1/S with 2.5 mM thymidine and then arrested in mitosis with 330 nM of nocodazole for 16 h before mitotic shake-off. Mitotic cells were then washed once by phosphate-buffered saline (Gibco), pelleted down, and chilled on ice. Lysis was performed by directly adding 2x Laemmli sample buffer (supplemented by 2-mercaptoethanol, Bio-Rad Laboratories) at a ratio of 1 µL per 0.1 mg of cell pellets and pipetting up and down. Lysates were boiled immediately afterward for 10 min and then chilled on ice. 8 µL of supernatant was loaded onto each lane of a 15-well, 0.75-mm SDS-PAGE mini gel.

Primary antibodies used included anti-BUBR1 (Bethyl Laboratories A300-995A, 1 : 1000), anti- MAD1 (GeneTex GTX109519, 1 : 2000), anti-CDC20 (Santa Cruz Biotechnology sc-13162, 1 : 200), anti-BUB3 (Sigma-Aldrich B7811, 1 : 500), anti-GAPDH (Proteintech 60004-1-Ig, 1 : 5000).

### RNA interference

Cells were transfected with siRNA in the morning. 2.5 mM of thymidine was added 8 h later and cells were incubated overnight. The next morning, cells were released from the thymidine block into fresh media. This sequence was then repeated once and cells were released into FluoroBrite™ DMEM (Gibco) supplemented with 9% (by volume) of FBS and 1× GlutaMAX and incubated for 6 h before the addition of mitotic drugs. Drugs were left to take effect for 1 h before imaging.

Sense-strand sequences and working concentrations of small interfering RNA duplexes (siRNAs) used in this study include the *SPC25* siRNA (5’- GCCUGCGAAGCAUUGUCCUACAUAA-3’, 40 nM ^53^), *BUBR1* siRNA (5’-GAUGGUGAAUUGUGGAAUA-3’, 40 nM ^54^), *BUB1* siRNA (5’-CGAAGAGUGAUCACGAUUU-3’, 40 nM ^55^), mNeonGreen siRNA (5’- GAGCUGAAGCACUCCAAGACA-3’, 40 nM) and the *ZW10* siRNA (5’-UGAUCAAUGUGCUGUUCAA-3’, 100 nM ^38^). The siRNAs against all five B56 isoforms were taken from the second pool in ^56^. Desalted siRNA duplexes modified by double-deoxythymidine overhangs at 3’-ends of both strands were synthesized by Sigma. The AllStar siRNA (QIAGEN) was used for the negative control. All siRNAs were transfected into the cells via Lipofectamine RNAiMAX (Invitrogen).

### Live-cell imaging

Cells were plated in a Nunc Lab-Tek II chambered cover glass (Thermo Scientific) or a 35-mm cover glass-bottomed dish (MatTek) and treated with drugs and/or siRNAs as described above. For imaging, the chambered cover glass or the cover glass-bottomed dish was loaded into a CU-501 temperature and gas control system (Live Cell Instrument). The sample holder was maintained at 37°C and ventilated by humidified 5% of CO2 and the objective was maintained at 37°C by a heating band.

MPS1sen-KT FRET and immunofluorescence experiments were performed on a Nikon Eclipse Ti-E/B inverted microscope, with a CFI Plan Apochromat Lambda 100×, 1.45 NA oil objective (Nikon). The microscope was equipped with an H117E1 motorized stage (Prior Scientific) and a NanoScanZ 100 piezo stage (Prior Scientific). A SPECTRA 5-LCR-XA Light Engine (Lumencor) served as the excitation light source. The 475 nm-centered band of excitation light was used for the green channel and the 575/30nm-filtered band of excitation light was used for the red channel. An ET-EGFP/mCherry filter cube (Chroma Technology) was used as the dichroic mirror, where the built-in emission filter on the cube has been removed. Emission light in the red channel was filtered by an ET632/60m (Chroma Technology). Emission light in the green channel was filtered by an ET525/50m (Chroma Technology). Emission filters were mounted on a high-speed filter wheel (Prior Scientific) positioned in the light path. Images were acquired with an iXon3 EMCCD camera (Andor Technology) operating in the conventional CCD mode. Signaling kinetochores in GSK923295-treated cells were identified by their polar positioning and the enrichment of localized SAC proteins on them.

The FRET measurement with MPS1sen-KT was performed on a Nikon Eclipse TiE/B inverted microscope, with a CFI Plan Apochromat VC 100×, 1.40 NA oil objective (Nikon). The microscope was equipped with an H117E1 motorized stage (Prior Scientific), a NanoScanZ 100 piezo stage (Prior Scientific), and an X-Light V2 L-FOV confocal unit with 60-µm pinholes (CrestOptics). A CELESTA Light Engine (Lumencor) served as the excitation laser source. The 477-nm line (at 25% power with an exposure time of 400ms for each frame) was used for both the green and the FRET channels and the 546-nm line (at 50% power with an exposure time of 400ms for each frame) was used for the red channel. A ZT488/543rpc (Chroma Technology) was used as the dichroic mirror. Emission through both the red and the FRET channels was filtered by an ET605/52m (Chroma Technology) while emission through the green channel was filtered by an ET525/36m (Chroma Technology). Images were acquired by a Prime 95B 25mm sCMOS camera (Teledyne Photometrics).

Time-lapse live-cell imaging related to the knockdown-rescue experiments was performed on an ImageXpress Nano Automated Imaging System (Molecular Devices). A SOLA Light Engine (Lumencor) served as the excitation light source. Cells were plated on 24-well cell imaging plates (black plate with treated glass bottom, Eppendorf) and treated with drugs and/or siRNAs accordingly. Input humidified 5% CO2 flow was maintained at around 19 psi and the environment chamber was maintained at 37°C. All SAC proteins tagged by mNeonGreen (BUBR1, BUB1, and MAD1) feature inhomogeneous distributions between the cytosol and the nucleus/nuclear envelope in the prophase (data not shown). In principle, they can indicate the accurate timing of the nuclear envelope breakdown (NEBD) in each cell during the time-lapse imaging. However, due to the resolution limit and for consistency, we determined mitotic duration mainly based on cell morphology (from rounding-up at the NEBD to elongation at the anaphase onset) from transmitted-light images. This was facilitated by the semi-automatic image analysis pipeline available at https://github.com/CreLox/IXNAnalysis ^48^.

### Immunofluorescence

Hela cells that exogenously expressing mNG-Bub1 were plated on a 12-mm coverslip and synchronized by thymidine (2.5 mM for 8 hours) followed by RO3306 treatment (15 nM for 20 hours). G2/M arrested cells were washed three times with PBS and released into media containing GSK923295 (236 nM) for 4 hours) to get arrested in Mitosis. For immunolabeling, the cells were pre-extracted with 0.5% Triton X-100 in PHEM (240 mM Pipes, 100 mM HEPES, 8 mM MgCl2 and 40 mM EGTA) and then fixed with 4% PFA for 10 min. The coverslips were washed three times with PHEM buffer and blocked for 30 min at room-temperature with 5% Donkey serum. Next, the coverslips were incubated in anti-Mad1 (1:1000, Sigma M80691), anti-Bub1 (1:500, Thermo A300-373A), or anti-BubR1 (1:1000, Thermo Scientific A300-373A) over night at 4°C. The next day, the coverslips were washed 4 times with PHEM containing 0.05% Tween-20 and incubated with appropriate Alexa Fluor 633-conjugated secondary antibodies (Goat anti-Mouse, Invitrogen A21052 or Goat anti-Rabbit, Invitrogen A21071) at a 1:10000 dilution for 45 min at room temperature in the dark. Following secondary antibody labeling, the coverslips were washed four times and mounted in an antifade solution (ProLong; Molecular Probes).

Immunofluorescence images were acquired on the spinning disk confocal microscope described above. For each field of view, 31 *z*-sections were acquired at 0.2-μm steps.

### Quantification of kinetochore-localized fluorescence signal

Quantification of the kinetochore-localized fluorescence signal was performed using a custom graphical user interface written in MATLAB. The methodology has been described previously ^11, 53^. Briefly, individual kinetochores were identified in the mCherry channel based on the Spc25- mCherry fluorescence. A 6×6-pixel box was drawn centered on the maximum intensity pixel in the in-focus mCherry plane, and the mNG intensity of all the pixels within this box was summed to obtain the mNG fluorescence signal for that kinetochore. The local background was defined as the median intensity value of the perimeter pixel intensities in a 10×10 pixel box concentric with the 6×6 pixel kinetochore box.

For immunofluorescence quantification involving both RPE-1 and HeLa cells, anti-ACA antibodies probed with Alexa Fluor 488-conjugated secondary antibodies were used to identify the centromeres. Alexa Fluor 633-conjugated secondary antibodies were used to visualize primary antibodies against the respective SAC proteins.

### Fluorescence correlation spectroscopy (FCS)

The total number of fluorophores in a homogeneous solution is *N*total := *N*A*cV*total, where *N*A is the Avogadro constant, *c* is the molar concentration of the fluorophore, and *V*total is the total volume of the solution. The probability that a specific fluorophore molecule is within the excitation volume *V*0(≪ *V*total) at any given time is *p*0 := *V*0*/V*total ≪ 1. For freely diffusive fluorophores in a diluted solution, whether or not a specific fluorophore is within the excitation volume is independent of each other. Thus, the number of fluorophores inside the excitation volume at any given time *N*0 has a binomial distribution *B*(*N*total*, p*0). Therefore, Under a fixed live-cell imaging setup [which includes the microscope (its alignment and the objective), the wavelength of the excitation light, the thickness of the coverslip (affecting the actual working distance), and the refractive index of the cytosol], *V*0 is fixed. Therefore, *G*0 is inversely proportional to the molar concentration of the fluorophore. The average number (or the variance of the number) of fluorophores inside the excitation volume observed over a long period should be close to the theoretical mean <*N*0> (or the theoretical variance *σ*^2^N0).

All FCS data were collected on an Alba v5 Laser Scanning Microscope (ISS), connected to an Olympus IX81 inverted microscope main body [equipped with a UPLSAPO60XW objective (1.2 NA, Olympus)]. A Fianium WL-SC-400-8 laser (NKT Photonics) with an acoustooptic tunable filter was used to generate excitation pulses at a wavelength of 488 nm and a frequency of around 20 MHz. Excitation light was further filtered by a Z405/488/561/635rpc quadband dichroic mirror (Chroma). Emission went through a 655DCSPXR short-pass dichroic mirror (Chroma) and an FF01-531/40-25 filter (Semrock) and was finally detected by an SPCM-AQRH-15 avalanche photodiode (Perkin Elmer). The time-correlated single photon counting module to register detected photon events to excitation pulses was SPC830 (Becker & Hickl). Data acquisition was facilitated by VistaVision (ISS). The excitation volume (*V*0) was calibrated by taking FCS data from TetraSpeck™ microspheres (0.1 µm, Invitrogen) of known concentrations.

### Statistical analysis

Statistical analysis was conducted using GraphPad Prism version 9.

## Acknowledgements

This work was supported by R35-GM126983 from the National Institute of General Medical Sciences to APJ. We thank Dr. Makayev for sharing the HeLa A-12 parental cell line, Dr. Geert Kops for kindly sharing the design details for the MPS1sen-KT construct, and Dr. Taylor for sharing anti-Bub1 antibodies. We also thank Dr. Iain Cheeseman and Dr. Mara Duncan for feedback on the manuscript and useful discussion.

## Supplementary figures

**Figure S1.**
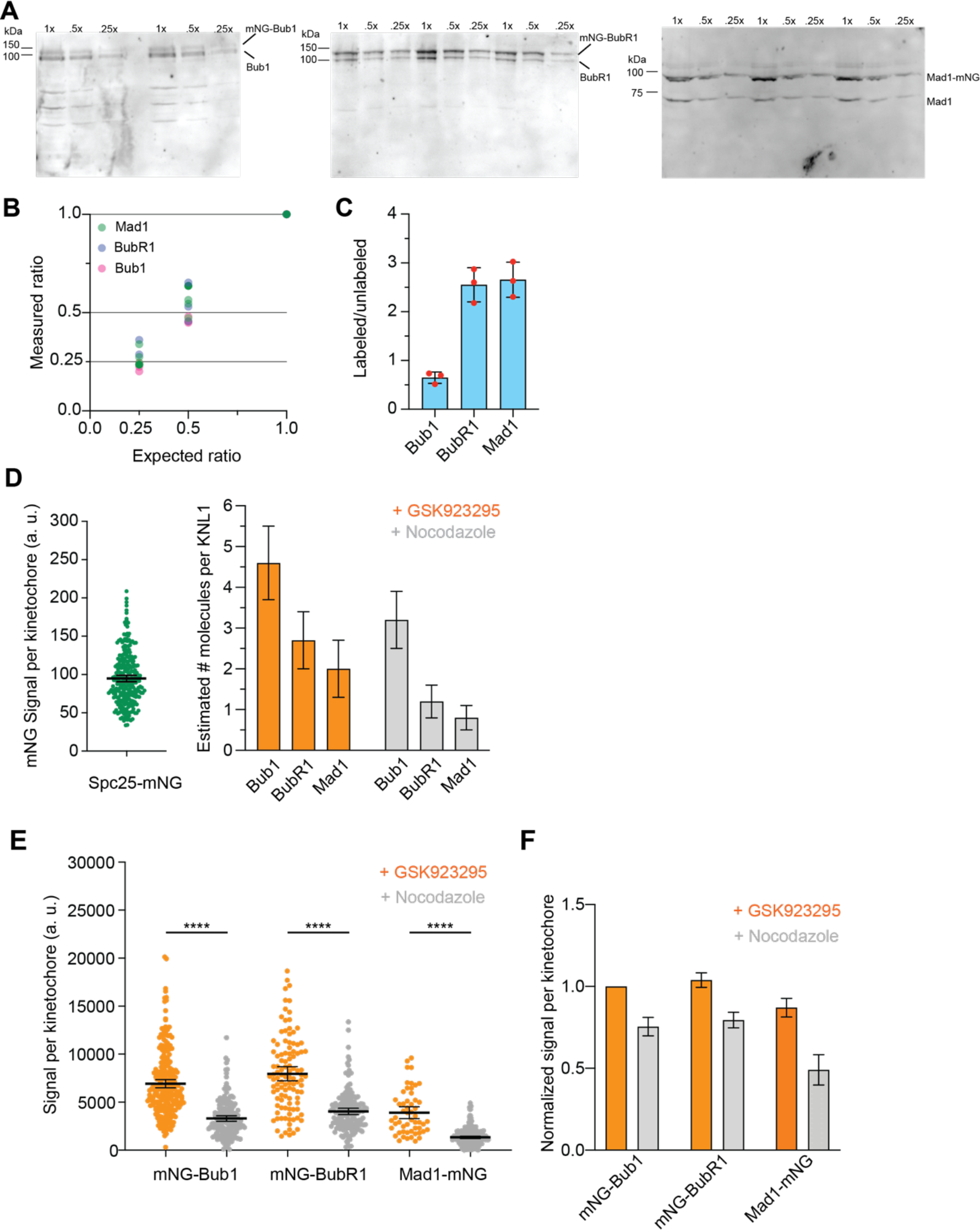
(related to Figure 1) Assessment of the relative amounts of mNG-labeled and unlabeled proteins in the three cell lines. **(A)** Immunoblot analysis of whole mitotic cell lysates of the three cell lines probing for the indicated proteins. Two out of the three technical repeats for the mNG-Bub1 cell line and all three technical repeats for the mNG-BubR1 and Mad1-mNG cell lines are displayed. Numbers at the top denote the lysate volume loaded to ascertain the linearity of the fluorescent secondary antibody signal. **(B)** Ratio metric analysis of serial dilutions confirms that band intensities are directly proportional to the amount of protein loaded. The band intensities were quantified using the ‘Gel Analyzer’ tool in Fiji. The ratio of labeled to unlabeled protein for each technical repeat was defined as the average of the ratios for the three ‘repeats’. (C) The ratio of the mNG-tagged to unlabeled species for the three proteins. Error bars display the standard deviation. (D) Left: the beeswarm plot displays the quantification of Spc25-mNG signal per kinetochore in images obtained under identical imaging conditions as in Figure 1C. The bar plot on the right displays the estimated number of molecules of the mNG-tagged SAC protein per Knl1 calculated by dividing the mean signal per kinetochore for each protein with the mean Spc25-mNG signal per kinetochore as the reference. Error bars represent mean ± SEM. **(E)** Left: Quantification of the fluorescence signal per signaling kinetochore in cells treated with siRNA against *ZW10* (n = 108, 242, 54 in GSK923295-treated cells and 166, 178, 219 in nocodazole-treated cells for Bub1, BubR1, and Mad1, respectively, pooled from 2 technical repeats). **(F)** Average signals from E are normalized using the mNG-Bub1 signal per kinetochore. Horizontal lines display the standard error on the mean ratio.

**Supplementary Figure S2.**
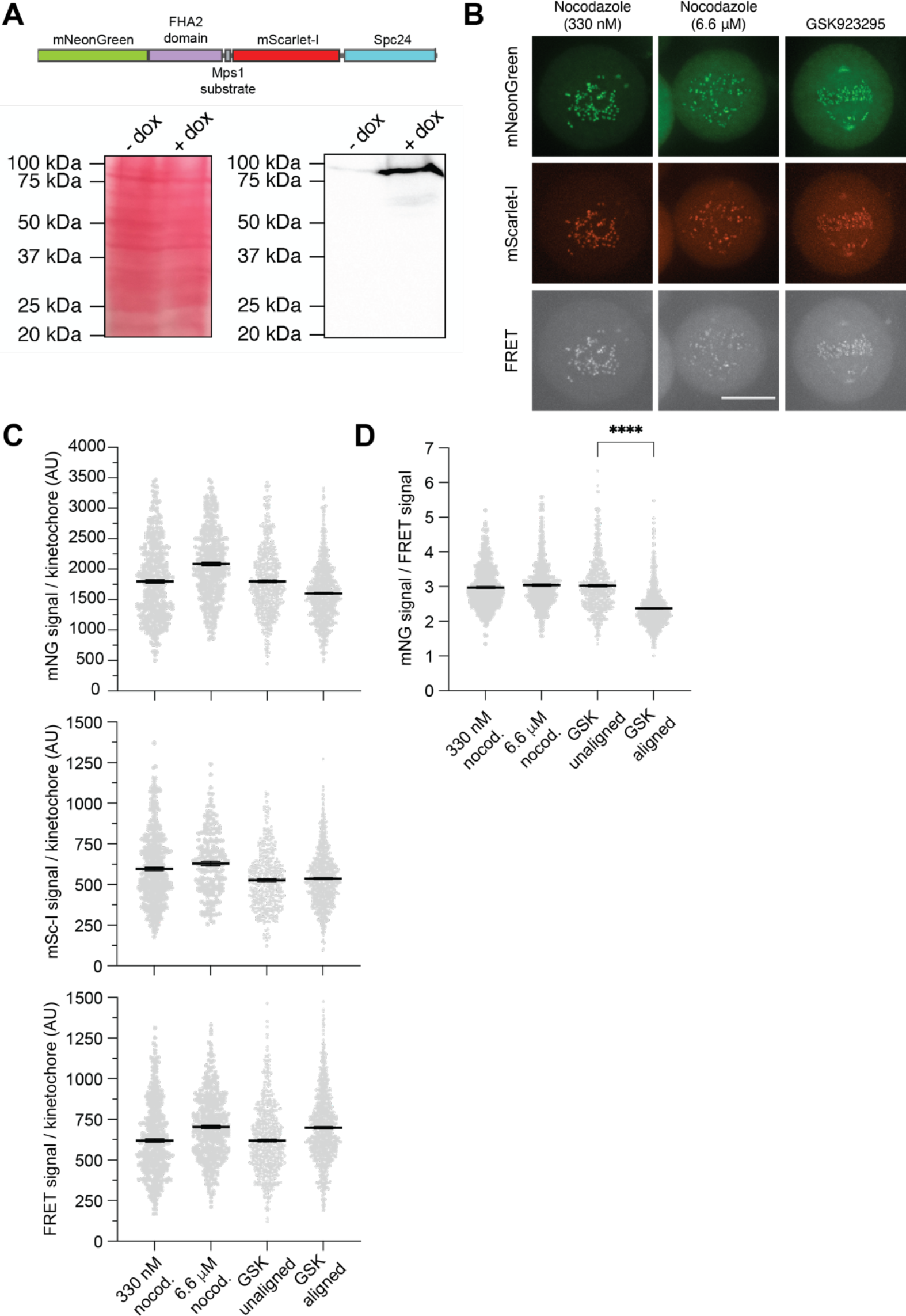
(related to Figure 1) Net phosphorylation at signaling kinetochores is the same in nocodazole- and GSK923295-treated cells. **(A)** Top: Schematic of the modified MPS1sen-KT ^34^ that uses the mNeonGreen and mScarlet-I as the FRET pair. Bottom: Using an antibody against DsRed2 (which detects mScarlet-I), we confirmed that MPS1sen-KT (with a theoretical molecular weight of 97.2 kDa) is expressed as a full-length protein with negligible partial degradation in HeLa A12 treated with doxycycline. A Ponceau S staining of the same blot before membrane blocking (left panel) serves as the loading and membrane transferring control. **(B)** Representative images of cells expressing MPS1sen-KT. The look-up table for each channel (row) is universal for different groups (column). Scale bar, 10 μm. **(C)** Raw fluorescence quantification at signaling kinetochores under various drug treatments in the green, red, and FRET channels. Each gray dot represents a single kinetochore measurement. Data from cells treated with 45 nM, 90 nM, and 200 nM GSK923295 were pooled together (the mean values are not significantly different from one another; data not shown) to simplify the presentation. Data were compiled from at least two independent experiments (more than 40 cells in total for each group). Mean values ± 95% confidence intervals are overlaid. The variation across different groups can be attributed to different FRET efficiencies (which we intend to quantify) and day-to-day variations in the doxycycline-induced expression of MPS1sen-KT. To cancel the effect of these variations, a normalized metric of the FRET efficiency (mNG/FRET) was used in this study. **(D)** A summary of a normalized FRET metric (mNeonGreen signal/FRET signal) in HeLa A12 cells treated with different drugs at various concentrations. Each gray dot represents a single kinetochore measurement. Data were compiled from at least two independent experiments (more than 40 cells in total for each group). As in C, Data from cells treated with 45 nM, 90 nM, and 200 nM GSK923295 were pooled together. Mean values ± 95% confidence intervals are overlaid. The Welch’s ANOVA test [*W*(DF*n*,DF*d*) = 1 .339(2.000,917.0), *P* = 0.2626] was performed for the first 3 columns (unaligned, signaling kinetochores). The unpaired t-test with Welch’s correction (*P*<0.0001) was performed to compare aligned non-signaling kinetochores with unaligned signaling kinetochores in GSK923295-treated cells.

**Figure S3.**
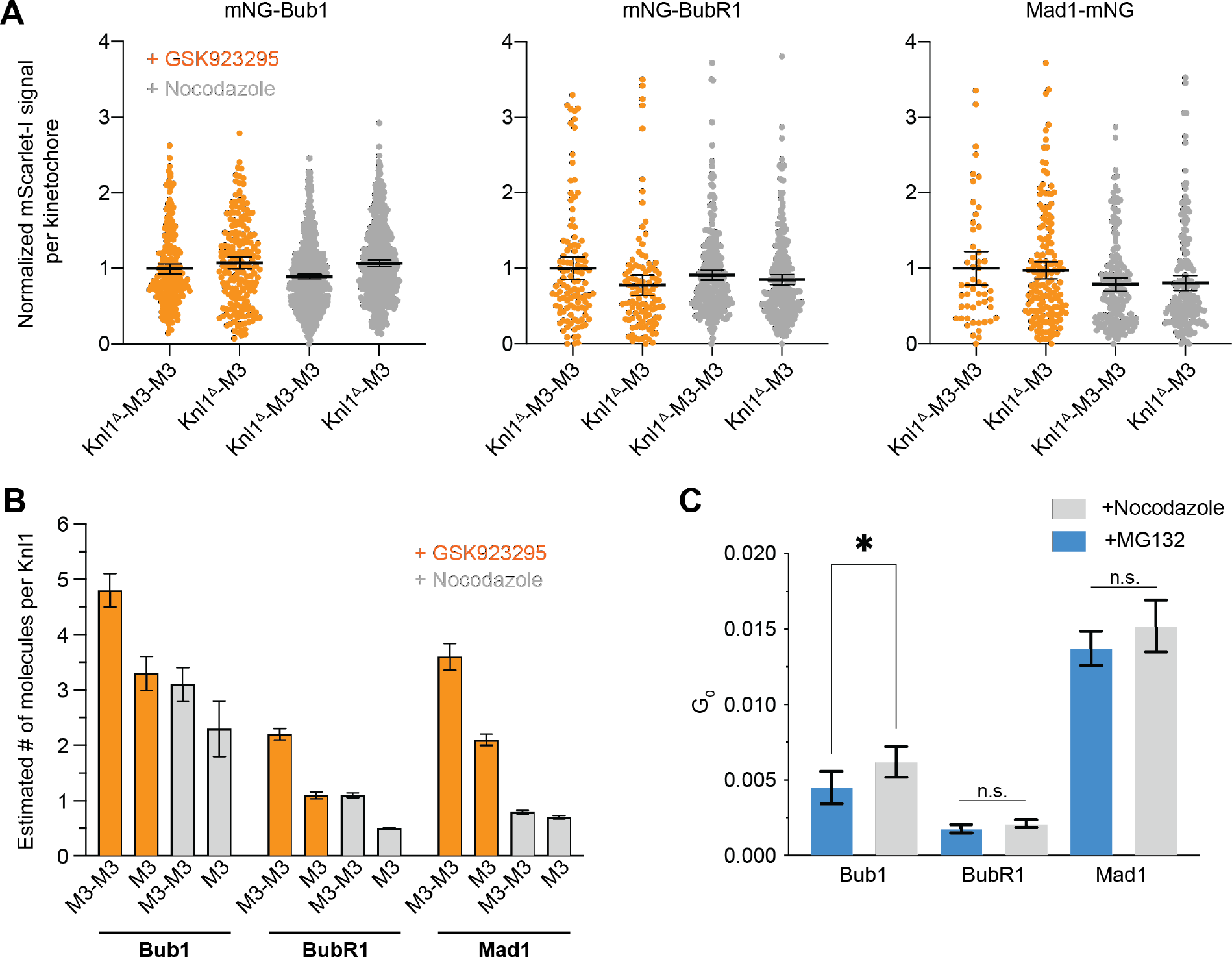
(related to Figure 2) Assessment of the kinetochore recruitment of mScarlet-I- Knl1^Δ^-M3-M3 and mScarlet-I-Knl1^Δ^-M3. **(A)** mScarlet-I fluorescence signal per kinetochore indicating the amount of recombinant Knl1 versions (noted by the X-axis labels). Minor differences in the incorporation of the recombinant Knl1 cannot explain the observed differences in SAC protein recruitment in each case. Note that the average value for the mScarlet-I-Knl1^Δ^-M3-M3 signal in GSK939295 in each data set was used as the normalization factor for each cell line. The numbers of data points are identical to those in Figure 2. **(B)** Estimation of the copy number of each SAC protein per Knl1 molecule using the Spc25-mNG fluorescence signal per kinetochore as the fiduciary standard (shown in Figure S1). Error bars indicate mean ± SEM. **(C)** *G*0 of the three proteins in metaphase-arrested (MG-132-treated) and nocodazole-treated cells as measured by Fluorescence Correlation Spectroscopy (FCS). This is the magnitude of the auto-correlation function for the fluorescence fluctuation data for 0-time delay. It is inversely proportional to the average number of fluorophores in the focal volume, i.e. the concentration of the fluorophore ^57^. Mean values under the two conditions compared using Welch’s *t*-test.

**Figure S4.**
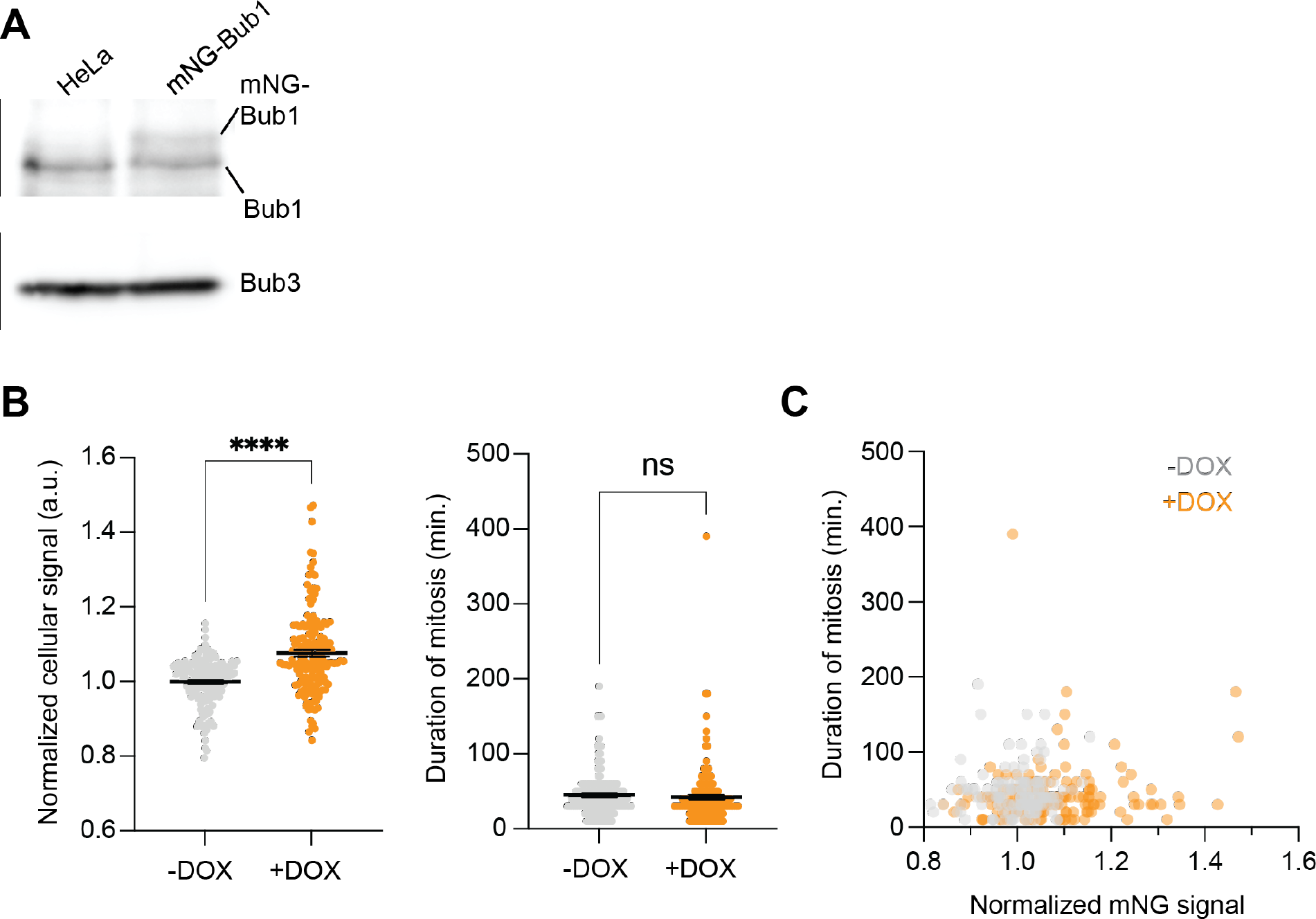
(related to Figure 3) mNG-Bub1 over-expression does not affect normal mitotic progression of HeLa A12 cells. **(A)** Immunoblot shows doxycycline-induced expression of exogenous mNG-Bub1. **(B)** Left: The average mNG signal in mitotic HeLa cells that are either untreated or treated with doxycycline to induce exogenous mNG-Bub1 expression (normalized to the average value of mNG fluorescence in untreated cells, which also represents the background signal). Right: the duration of mitosis for the same cells. (n = 155 and 154 for -dox and +dox respectively pooled from 2 technical repeats). The horizontal lines indicate mean ± SEM. Welch’s *t*-test was used to compare the sample means. **(C)** Scatterplot displays the correlation between the cellular mNG signal (normalized as in B) and the duration of mitosis (Pearson’s correlation coefficient = 0.05 and 0.13 with *p* = 0.5 and 0.1 for -Dox and +Dox respectively).

## Supplementary Videos

**Video S1 –** Time-lapse imaging of HeLa cells treated with 25 nM GSK923295. Time - hr:min.

**Video S2 -** Time-lapse imaging of HeLa cells treated with 25 nM GSK923295 and 1 μg/ml of Doxycycline to exogenously over-express mNG-Bub1. Time - hr:min.

**Video S3** - Time-lapse imaging of HeLa cells treated with 330 nM Nocodazole. Time - hr:min.

**Video S4 -** Time-lapse imaging of HeLa cells treated with 330 nM Nocodazole and 1 μg/ml of Doxycycline to exogenously over-express mNG-Bub1. Time - hr:min.

**Video S5 –** Time-lapse imaging of genome-edited HeLa cells heterozygous for mNG-Bub1 and exogenously expressing H2B-mCherry. Cells were released from a G1/S block into media containing 236 nM GSK923295. Time - hr:min.

**Video S6 –** Time-lapse imaging of genome-edited HeLa cells heterozygous for mNG-Bub1 and exogenously expressing H2B-mCherry. Cells were treated with siRNA targeting mNG to partially deplete BubR1 and released from a G1/S block into media containing 236 nM GSK923295. Time - hr:min.

